# Nomadic Lifestyle of *Lactobacillus plantarum* Revealed by Comparative Genomics of 54 strains Isolated from Different Niches

**DOI:** 10.1101/043117

**Authors:** Maria Elena Martino, Jumamurat R. Bayjanov, Brian E. Caffrey, Michiel Wels, Pauline Joncour, Sandrine Hughes, Benjamin Gillet, Michiel Kleerebezem, Sacha A.F.T. van Hijum, François Leulier

## Abstract

The ability of many bacteria to adapt to diverse environmental conditions is well known. Recent research has linked the process of bacterial adaptation to a niche to changes in the genome content and size, showing that many bacterial genomes reflect the constraints imposed by their habitat. However, some highly versatile bacteria are found in diverse niches that almost share nothing in common. *Lactobacillus plantarum* is a lactic acid bacterium that is found in a large variety of niches. With the aim of unravelling the link between genome evolution and ecological versatility of *L. plantarum,* we analysed the genomes of 54 *L. plantarum* strains isolated from different environments. Phylogenomic analyses coupled with the study of genetic functional divergence and gene-trait matching analysis revealed a mixed distribution of the strains, which was uncoupled from their environmental origin. Our findings demonstrate the high complexity of *L. plantarum* evolution, revealing the absence of specific genomic signatures marking adaptations of this species towards the diverse habitats it is associated with. This suggests fundamentally similar and parallel trends of genome evolution in *L. plantarum,* which occur in a manner that is apparently uncoupled from ecological constraint and reflects the nomadic lifestyle of this species.

## Introduction

Lactic acid bacteria (LAB) are a group of Gram-positive acid-tolerant bacteria that occupy a wide range of niches. The largest and most diverse genus of LAB is *Lactobacillus*, which includes more than 200 species (http://www.bacterio.net/lactobacillus). The environmental distribution of *Lactobacillus* species is highly variable; some species are exclusively found in specific habitats (e.g., *Lactobacillus helveticus* and *Lactobacillus delbrueckii* ssp. *bulgaricus* in diary products, *Lactobacillus johnsonii* and *Lactobacillus gasseri* in vertebrate gastrointestinal tracts) and other species, such as *Lactobacillus plantarum* and *Lactobacillus casei*, are encountered in a variety of different environments ^1,2^.

*L. plantarum* is an extremely versatile LAB that has been isolated from a variety of niches, such as plants, the gastro-intestinal tracts of human, animals, as well as food materials, like meat, fish, vegetables and raw or fermented dairy products ^1,3^ Its wide industrial utility and potent impact on animal phyisiology have made *L. plantarum* an organism of significant interest to the scientific community. It is one of the best-characterised vegetal associated bacteria that transform a multitude of plant-derived raw materials through fermentation ^4,5^ Also, as some *L. plantarum* strains are naturally-ocurring human commensals, they have been marketed as probiotics ^1,6,7^ and their potential beneficial effects on human or animal health have been recently reported as being able to promote juvenile growth in both *Drosophila melanogaster* and mouse in the presence of nutritional challenges ^1,7-13^.

Strains of different *Lactobacillus* species have been reported to adapt to defined environments by genome specialization driving niche-specific fitness. For example, *L. reuteri* strains isolated from different vertebrate intestinal tracts display host-adapted genome evolution paths ^14-18^; *L. paracasei* and *L. delbrueckii* strains adapt specifically to dairy environments through a process characterized by genome decay ^19-21^. *L. plantarum* is a generalist species that encompasses highly diverse strains whose evolutionary relatedness is not clear. Several studies have depicted the genetic diversity of *L. plantarum* through different phenotypic ^1,2^ and genotypic approaches, such as AFLP, RAPD ^1,3,15,18,22-24^, multi-locus sequence typing (MLST) ^4,20,21^, and microarray-based comparative genome hybridization (CGH) ^1,7^. However, despite these insights, whether a potential connection exists between a specific genomic background and a defined source of isolation remains an open question. High-throughput genomic approaches provide a deep understanding of the evolution and ecology of microorganisms. They help to assign putative adaptive features as well as putative functional roles to a given species/strain in an ecosystem. With the aim of gaining more insights into the functional capabilities and differences of *L. plantarum*, we sequenced the genomes of 43 *L. plantarum* strains that were isolated from a variety of food environments (such as fermented vegetables, dairy products, fruit and meat) and two natural animal hosts (human and *Drosophila melanogaster*). We next compared these 43 genomes to the existing genomes of 11 *L. plantarum* strains that were publicly available at the time we started our analyses (6 complete and 5 draft genomes). Our analysis therefore includes a wide-range of diverse strains, which enables the in depth analysis of *L. plantarum* phylogenomics. We now report the comparative analysis of these genomes with the main objective of exploring the potential link between the intra-species genetic variability and niche of isolation.

## Results

### Comparative analysis of 54 *L. plantarum* strains

To broadly investigate the genomic diversity of *L. plantarum* species, we chose 54 *L. plantarum* strains whose origins encompassed a large spectrum of ecological niches: 17 strains isolated from fermented fruits and vegetables, 11 strains from human origin (oral cavity, urine and intestinal tract, faeces and one putatively from the spinal fluid), 7 strains isolated from silage, 6 strains of dairy origin, 6 strains isolated from meat products and 6 strains isolated from adult *D. melanogaster’s* midgut (Table 1). The draft genome sequences of the 43 *L. plantarum* strains were analysed together with 11 publicly available *L. plantarum* genomes (Table 1) and aligned to the *L. plantarum* WCFS1 reference genome. In order to share all the results of the comparative analyses conducted on these genomes, we have created an online database that presents detailed information concerning the strain-specific gene content relative to the overall pan-genome of the 54 *L. plantarum* strains, including a comparative analysis of functionalities encoded in the individual genomes. The database is available at http://igfl.ens-lyon.fr/equipes/f.-leulier-functional-genomics-of-host-intestinal-bacteria-interactions/l-plantarum/. The sizes of the sequenced genomes range from 3 to 3.6 Mb, and encompassed a set of 1957 Orthologous Groups (OGs) that are shared by all *L. plantarum* strains (core genome). The pangenome size of the 54 genomes amounts to 7107 OGs (Table D1 available at http://igfl.ens-lyon.fr/equipes/f.-leulier-functional-genomics-of-host-intestinal-bacteria-interactions/l-plantarum/), which is consistent with what has been previously reported for many *Lactobacillus* species ^1,7,17,19,25^. Nevertheless, the pan-genome did not appear to approach saturation with the current strain collection, implying that the genetic repertoire of *L. plantarum* exceeds the current pan-genome estimate (Supplementary Fig. 1). The core genome of 1957 OGs (Table D2 available at http://igfl.ens-lyon.fr/equipes/f.-leulier-functional-genomics-of-host-intestinal-bacteria-interactions/l-plantarum/) covers 66% of *L. plantarum* WCFS1 annotated genes, and is of a similar size as has been reported for other *Lactobacillus* species ^1,15,21,26^ Moreover, the core-genome size is in good agreement with that obtained by microarray-based CGH analysis of *L. plantarum* strains that predicted 2049 core genes ^1,25^, and used 20 strains, which could explain the somewhat larger size of the core-genome estimate. The *L. plantarum* core genome contains the anticipated shared genetic repertoire involved in replication, transcription, and translation, as well as genes involved in energy production and amino acid‐ and carbohydrate-transport and metabolism (Supplementary Fig. 2). The variome in the *L. plantarum* collection contains 5150 OGs, among which 4500 OGs appeared to be scaffolded in the chromosome (Table D3 available at http://igfl.ens-lyon.fr/equipes/f.-leulier-functional-genomics-of-host-intestinal-bacteria-interactions/l-plantarum/). This estimate may be slightly high, since a fraction of the OGs represent fragments of mobile element (e.g. transposases of the IS elements) that can be part of scaffolds, or part of independently replicating plasmids. We estimate that there are 2686 to 3253 OGs per strain (Table 1), of which 2600-3000 OGs are on contigs in scaffolded chromosomes and 0-300 are non-scaffolded and may be encoded by plasmids. Most of the sequenced *L. plantarum* strains appeared to have several plasmids, with the exception of 6 strains (CNW10, NIZO2264, NIZO2855, NIZO2877, NC8 and JDM1). The list of genes that appear to be encoded by plasmids is reported in Table D4 available at http://igfl.ens-lyon.fr/equipes/f.-leulier-functional-genomics-of-host-intestinal-bacteria-interactions/l-plantarum/. Our comparative genome analysis identified 4137 novel OGs that are not present in the reference genome WCFS1 (Supplementary Table 1).

**Table 1.**
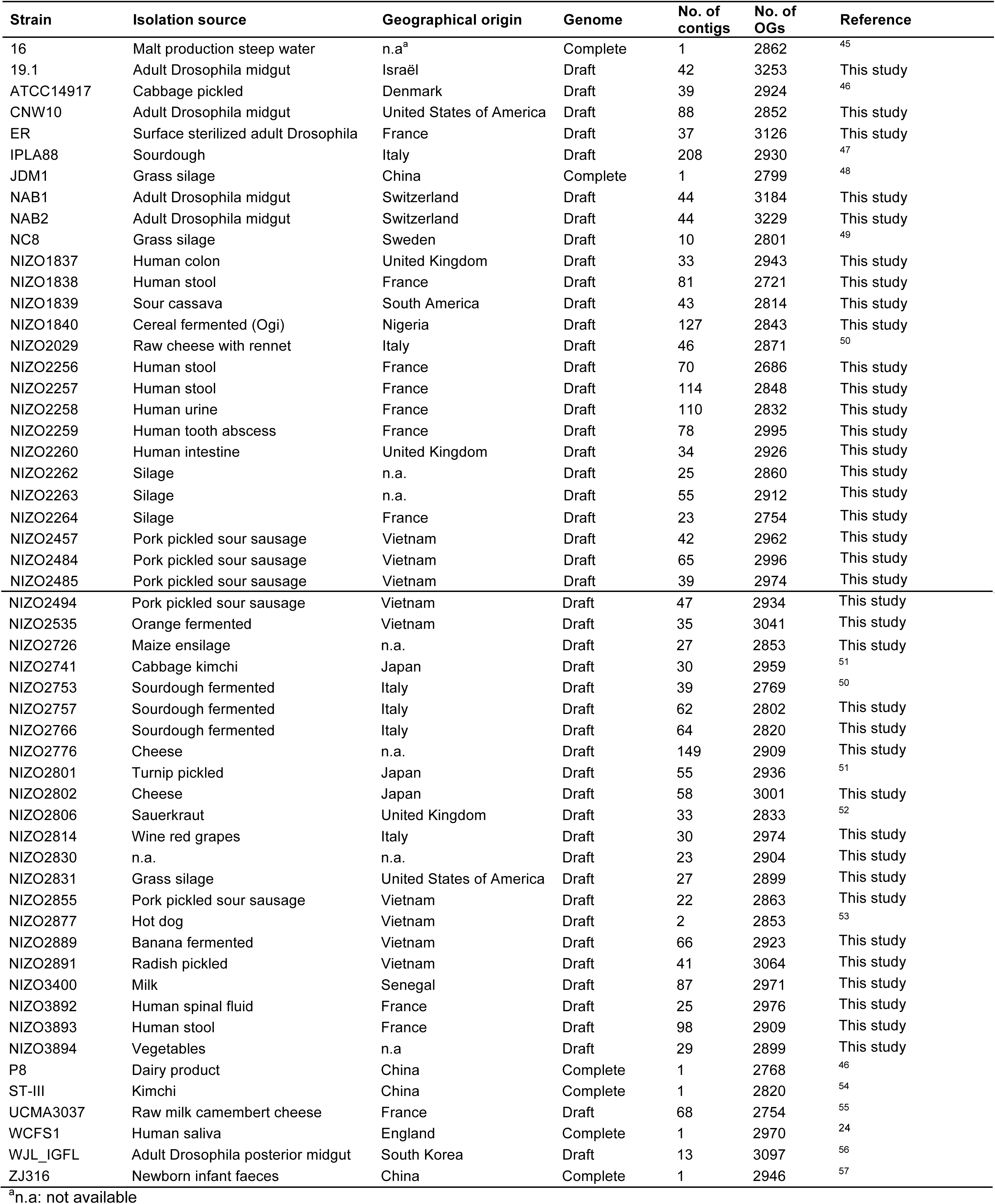
*L. plantarum* strains characterized in this study.

We evaluated the degree of gene content variation among the 54 genomes by analysing the presence/absence of the OGs encoding proteins associated to central cellular processes and/or molecular functions. As reported by Molenaar and colleagues ^7^, the highest genetic conservation in *L. plantarum* strains is observed for OGs involved in energy metabolism, or biosynthesis or degradation of cellular structural components, such as nucleotides, proteins and lipids (for further details see also section “Variable Regions” available at http://igfl.ens-lyon.fr/equipes/f.-leulier-functional-genomics-of-host-intestinal-bacteria-interactions/l-plantarum/database-for-comparative-genomics-of-54-l-plantarum-strains-2-7).

Despite the high conservation of these biosynthetic pathways, *L. plantarum* shows high inter-strain genetic variability. Consistent with previous studies, large variable regions exist among the 54 *L. plantarum* genomes ^1,7,25^ The most variable regions include genes involved in exopolysaccharide (EPS) biosynthesis, restriction-modification, sugar-importing phosphotransferase (PTS) systems and other transport functions, sugar metabolism, bacteriocin production, as well as the notoriously variable functions associated with prophages, insertion-sequence (IS) elements and transposases, which is in agreement with previous studies ^1,7,25^ (for further details see also section “Variable Regions” available at http://igfl.ens-lyon.fr/equipes/f.-leulier-functional-genomics-of-host-intestinal-bacteria-interactions/l-plantarum/database-for-comparative-genomics-of-54-l-plantarum-strains-2-7).

### Gene content analysis does not reveal specialization of *L. plantarum* for a specific environment

The process of niche adaptation may include events of gene gain and/or loss, or genome decay. Genes whose functions are dispensable for a strain’s fitness in a particular environment can be lost during niche adaptation. We therefore analysed the potential link between strain origin and their gene content. We first sorted strains by comparing the total number of OGs they encode (Supplementary Fig. 3), and found no strain grouping dependent on the origin of isolation, although it is quite remarkable that 5 out of the 6 strains isolated from *Drosophila melanogaster’s* midgut (19.1, ER, NAB1, WJL_IGFL, NAB2) encode the highest number of OGs. Next, we generated a dendrogram of the 54 *L. plantarum* strains based on the presence/absence of each OG in their pan-genome (Fig. 1). Two major groups could be distinguished: strains isolated from dairy products belong to group A, strains isolated from meat products and *Drosophila melanogaster* cluster in group B, while strains of human and plant origin are spread across the two groups. Notably, in each group, strains isolated from the same niche do not appear to be closely related (i.e., within the same subcluster), indicating that gene distribution poorly reflects strain origin.

**Figure 1.**
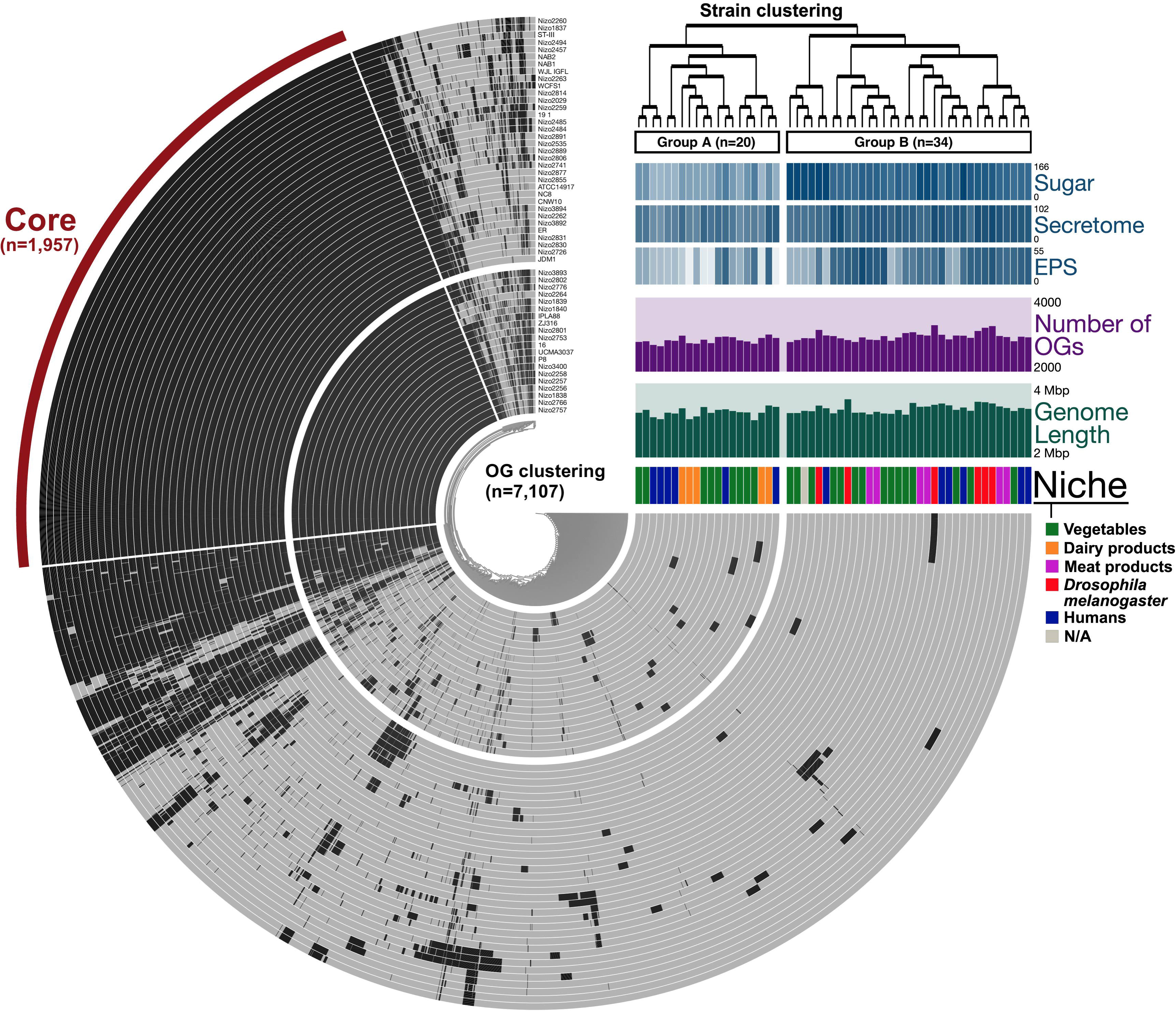
OG distribution across *L. plantarum*. The center of the figure shows the hierarchical clustering of OGs based on their presence/absence. Each ring represents a *L. plantarum* strain and each layer displays the OG distribution. Black colour stands for OG presence, whereas grey colour represents OG absence. The red bar groups the OGs belonging to the core genome. The top right panel reports additional metadata information about the dataset. The clustering present on the top right refers to strain grouping based on OG distribution. The occurrence of genes belonging to the 3 most variable genetic regions (exopolysaccharide biosynthesis (EPS), sugar cassettes, and secretome) is shown. The colour intensity of each bar reflects the increasing number of OGs present. Each strain (layer) is labelled depending on the source of isolation. The figure has been generated using Anvi’o software ^44^.

Next, using the PhenoLink tool ^27^, we performed a Gene Trait Matching (GTM) analysis, a statistical method based on the Random Forest algorithm that is used to correlate gene presence and absence to phenotypic features ^26,28,29^. We used the strains’ origins and OG presence/absence as inputs for the GTM. Among the most important associations, we identified groups of genes whose presence or absence associated with the origins isolated from vegetables, meats, humans and *Drosophila melanogaster*. No gene association could be pinpointed for the dairy origin (Supplementary Table 2). However, when we defined the niche isolation signature as genes present in strains isolated from a niche but mostly absent in all the others strains or *vice versa*, we found few genes: mostly hypothetical proteins or genes related to conjugation in the *Drosophila melanogaster* strains, bear such a robust niche isolation signature. However, currently it is difficult to assign specific functions to these genes in terms of niche adaptation. Therefore, these observations indicate that gene presence/absence is a poor indicator of origin for the considered *L. plantarum* strains.

### Phylogeny and analysis of allelic and functional divergence in *L. plantarum* confirm the absence of niche-specialization

Niche adaptation may not just depend on gene absence or presence, but can also be the result of accumulation of adaptive specific allelic variations of conserved genes. To gain insights into the potential role of such allelic variation in niche specialization and to reveal the evolutionary relationships of the *L. plantarum* strains studied, we performed a phylogenetic analysis of the core genome of the 54 strains (Fig. 2). The core genome phylogeny clearly shows a mixed distribution of strains isolated from similar sources. Thus, no phylogenetic cluster related to a defined niche appears to be present, but pairs of strains sharing more similar core genomic OGs were identified (NIZO2891 and NIZO2535, NIZO2484 and NIZO2484, NIZO2877 and NIZO2855, NIZO2260 and NIZO1837, NIZO2494 and NIZO2457, NIZO2831 and NIZO2726, NIZO2262 and NIZO3894). These strains are not clonal but share a highly similar set of core genome associated OGs; however, they also have the same geographical origin (Table 1), therefore their phylogenetic relatedness cannot be solely attributed to a process of niche adaptation.

**Figure 2.**
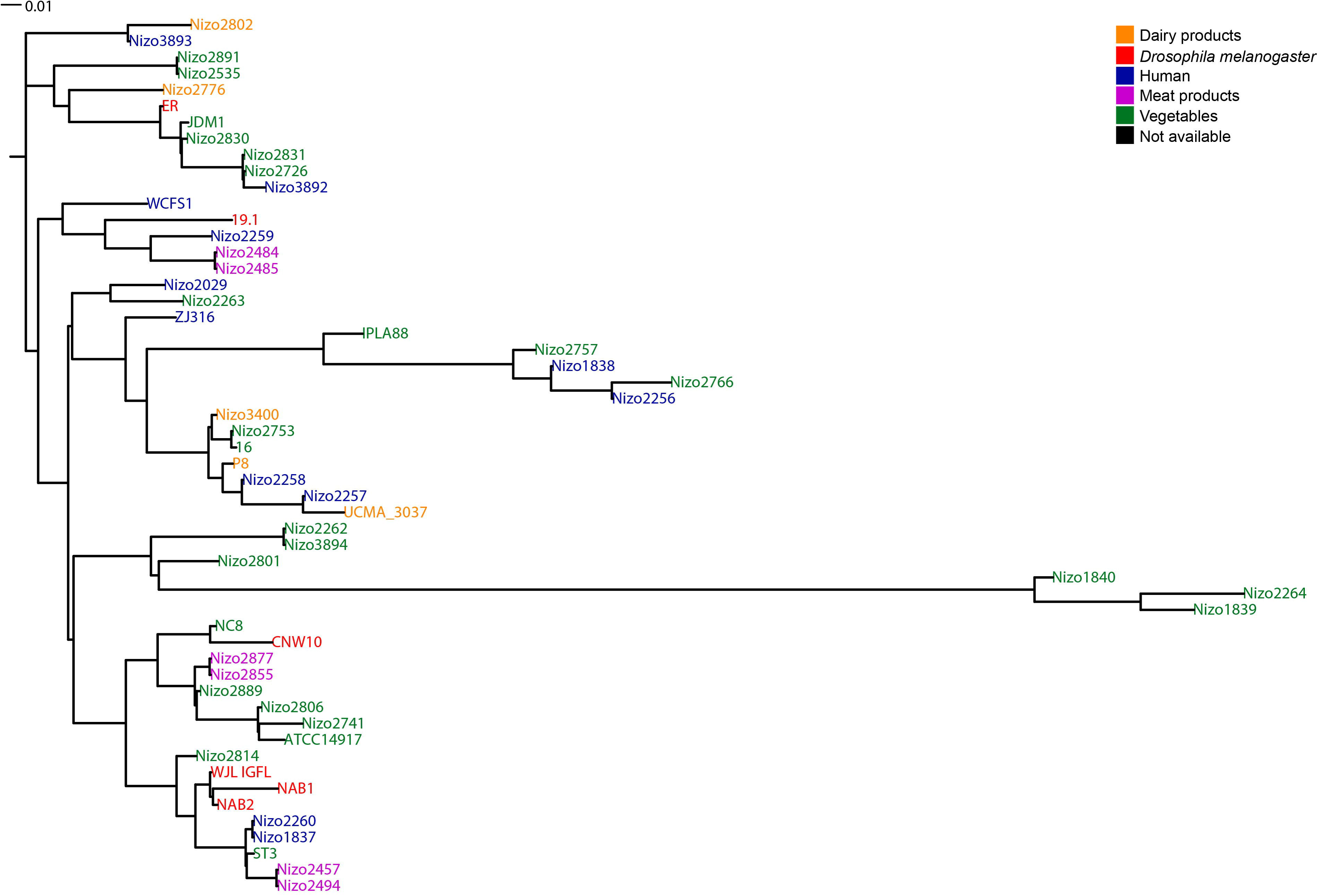
Phylogeny of *L. plantarum*. The 54 *L. plantarum* strains were clustered based on the nucleotide sequence of the core genome.

To further investigate the evolutionary history of *L. plantarum* strains and find potential niche-adaptation signatures, we tested if the phylogenetic relatedness of the strains is coupled to any functional divergence, i.e., the process by which new genes and functions originate through modification of existing ones. To probe for functional divergence, we employed the software tool developed by Caffrey et al. ^30^ to pinpoint enrichment or impoverishment of amino-acid replacements within the predicted core-proteome; enrichment may indicate specialization, wherease impoverishment signifies fixation. Subsequently, correlating such functional divergence detected in the individual proteins to the source of isolation can reveal whether amino-acid replacement(s) may have influenced the evolution of new functions within a group of strains originating from the same source. Importantly, this analysis takes into account the potential occurrence of horizontal gene transfer, assuming that the phylogenies of individual proteins may differ from the whole genome phylogeny, and provides a different representation of the evolutionary relationships among the strains. As a result, we found that 80% of the core proteome was either impoverished or not enriched for functional divergence among strains compared to the background (Fig. 3a). The impoverished (30%) Clusters of Orthologous Groups (COGs) are almost all related to genome information storage and processing such as DNA replication, recombination and repair, transcription, translation, ribosomal structure and biogenesis, protein turnover and chaperones. Caffrey et al. previously made similar observations based on the analysis of 750 complete bacterial proteomes ^30^ but their study also included COGs related to transport and metabolism of carbohydrates and nucleotides. The remaining 20% of COGs were found to be enriched in functional divergence (Fig. 3a), they include cellular defence mechanisms (V), cell motility (N), secondary metabolites biosynthesis, transport and catabolism (Q) and intracellular trafficking, secretion and vesicular transport (U) (Supplementary Table 3). However, the enrichment of functional divergence in these COGs is represented in a very limited number of strains (C: 5, U: 2, Q: 1, N: 1) of varying origin of isolation (Fig. 3b), implying that functional enrichment and source of isolation are not correlated. This conclusion was further supported by clustering the strains based on the functional divergence of each COG group, which is important to verify whether the strains eventually cluster according to their source of isolation (Fig. 3b). Taken together, these results indicate that the functional divergence of the *L. plantarum* strains is not correlated with their source of isolation.

**Figure 3.**
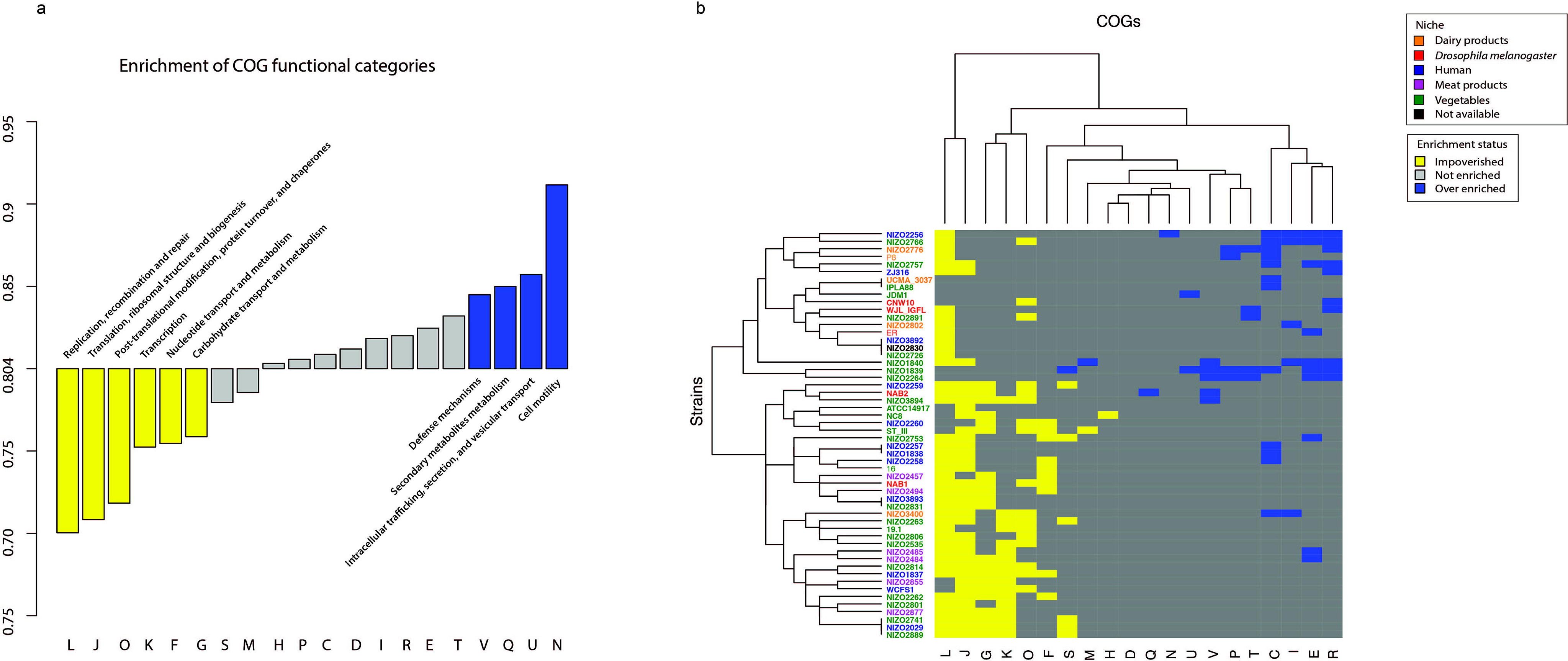
Analysis of functional divergence in *L. plantarum*. (a) Functional divergence of clusters of orthologous groups (COGs) was classified into three categories; impoverished (yellow, 6 groups), enriched (blue, 4 groups), or neither impoverished nor enriched (grey, 10 groups). (b) Heatmap showing the hierarchical clustering of the main patterns of functional divergence in the 54 *L. plantarum* strains (tree on the left hand side, and strain identifiers on the right hand side), as well as COG category clustering (shown on the top).

### The analysis of highly variable regions in *L. plantarum* supports the lack of niche-adaptation processes

Thus far, the analysis of the gene content, the phylogenetic study of the core genome and the analysis of the proteome for functional divergence failed to identify any niche specialization signatures in the genome of the 54 *L. plantarum* strains. However, the phylogenetic analysis reported above does not take into account the accessory genes (variome) of the 54 *L. plantarum* strains. In addition, when we clustered pan-genome OG distribution based on the presence/absence of OGs belonging to the most variable functionalities (secretome, sugar metabolism and EPS), we found a potential signature of clustering into Group A or B (Fig. 1). Therefore, the core genome phylogeny depicts the evolution of the genes that are presumably necessary for all strains, but the distribution pattern of the variome might reflect a specific evolutionary trajectory taken by the respective strains. Therefore, we decided to focus on the variable functional categories and analyse each of them separately to evaluate whether the OG content of a particular functional category may predict the strains’ origins. We performed independent hierarchical clustering of 3 of the functional categories that displayed the highest variability in terms of OG presence and absence among the 54 *L. plantarum* strains (i.e., EPS, sugar utilization cassettes, and secretome) (Fig. 4). In addition, we performed independent allelic divergence analyses (Supplementary Figs 4-6) within the shared gene sequences of each category.

**Figure 4.**
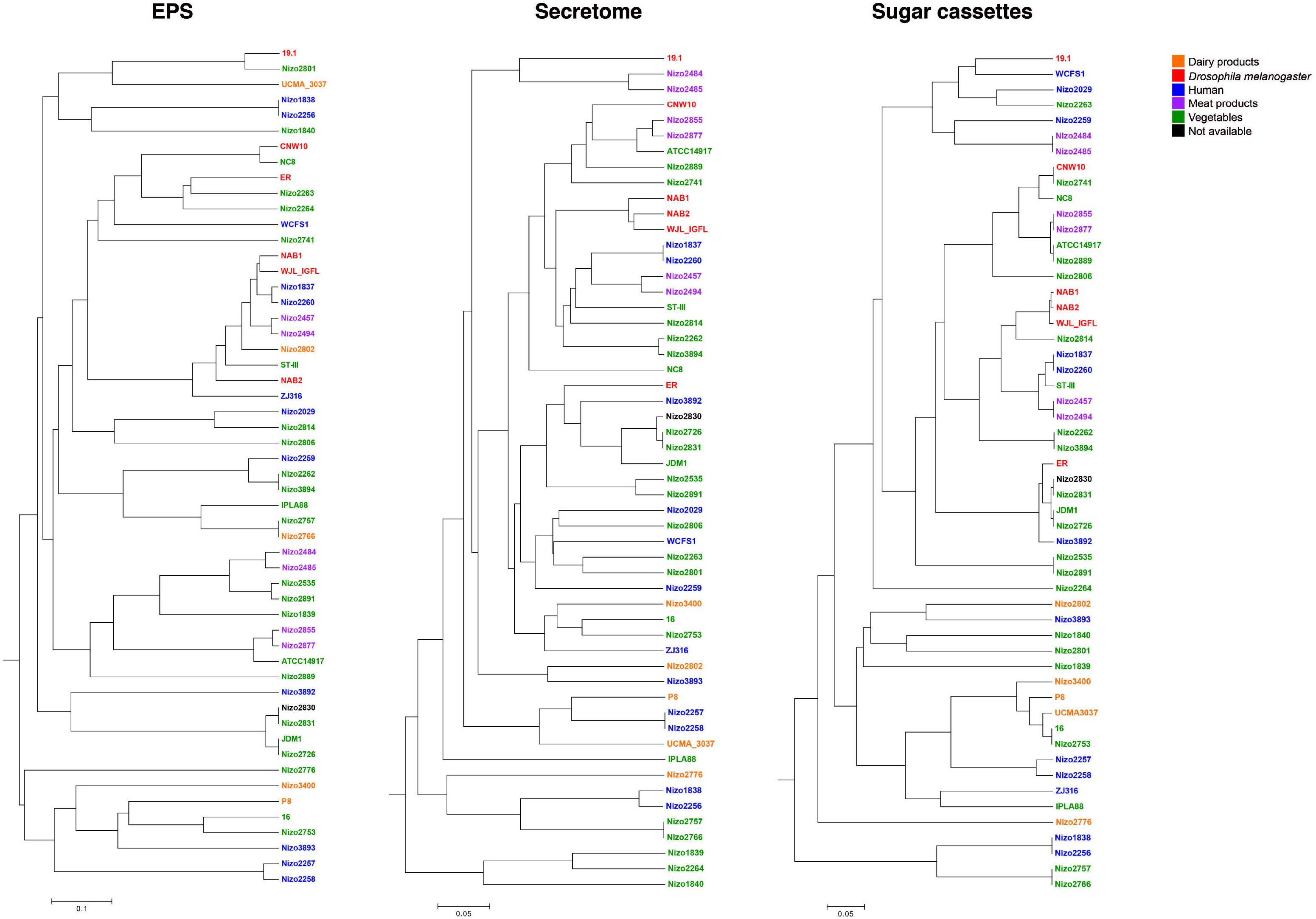
Hierarchical clustering of the 54 *L. plantarum* strains based on the most variable regions. The 54 *L. plantarum* strains were clustered based on the presence/absence of the genes belonging to the EPS, sugar metabolism and secretome categories.

#### EPS/CPS biosynthesis genes

The gene clusters involved in the EPS/CPS biosynthesis vary considerably in size, composition, sequence and gene order across the 54 *L. plantarum* strains. A detailed description and comparative analysis of these regions is reported at http://igfl.ens-lyon.fr/equipes/f.-leulier-functional-genomics-of-host-intestinal-bacteria-interactions/l-plantarum-on-line-material/eps-text-description.

The distribution and allelic divergence analysis of the EPS/CPS-assigned OGs (Fig. 4, Supplementary Fig. 5, respectively) are significantly correlated (p-value = 9.99001e-04), revealing that the clustering of the strains on basis of the allelic divergence of the OGs in this category is strongly driven by the presence/absence distribution of these OGs. However, unlike for what has been observed for *L. paracasei*^21^, no correlation was found between the source of isolation and the presence of specific (subsets of) EPS/CPS-assigned OGs (Fig. 4). The strains with the fewest EPS/CPS-assigned OGs as well as those with the most EPS/CPS-assigned OGs are of various origins (human, dairy products and vegetables) (Table D9 available at http://igfl.ens-lyon.fr/equipes/f.-leulier-functional-genomics-of-host-intestinal-bacteria-interactions/l-plantarum/database-for-comparative-genomics-of-54-l-plantarum-strains-7-7). Moreover, strains isolated from the same niche appear to be scattered across the phylogenetic tree (Supplementary Fig. 4), and strains that contain a similar set of EPS/CPS-assigned OGs were isolated from different niches.

#### Secretome

The OGs assigned to the secretome were sorted and categorized into functional categories (Table D10 – Table D11 available at http://igfl.ens-lyon.fr/equipes/f.-leulier-functional-genomics-of-host-intestinal-bacteria-interactions/l-plantarum/database-for-comparative-genomics-of-54-l-plantarum-strains-7-7). Consistent with previous studies ^15,21,31^, the most variable sub-categories of the secretome included cell-surface complexes (*csc*), ATP-binding cassette (ABC) transporters and bacteriocins.The distribution of secretome-assigned OGs among the 54 *L. plantarum* strains revealed no correlation with the origin of isolation of the strains (Fig. 4). Analogous to what was observed for the allelic divergence analysis of the EPS/CPS assigned OGs, the allelic divergence analysis of the secretome-assigned OGs appeared to be quite strongly driven by the presence/absence of these genes among the strains. However, the refinement of this clustering by the allelic divergence did not reveal an association to the origin of isolation (Supplementary Fig. 5).

#### Sugar metabolism

Sugar metabolism genes are highly variably distributed among lactic acid bacteria (LAB) ^1,2,7,21^ The presence or absence of these genes presumably reflects adaptation to the availability of substrates for growth in different niches. Previous comparative studies conducted on other LAB species showed a strong correlation between the source of isolation and sugar metabolism ^15,31^. *L. plantarum* has been reported to contain a highly variable repertoire of genes related to sugar import and utilization, which appears to cluster in a so-called sugar utilization island of the genome ^1,2,7,22^. However, to date, no clear relationship has been deduced between the sugar utilization repertoire of different strains of *L. plantarum* and their source of isolation. The genomic analysis of the 54 *L. plantarum* strains confirmed the high plasticity and diversity of the genetic repertoire related to sugar metabolism in this species. The distribution of all OGs assigned to be involved in sugar metabolism in the 54 *L. plantarum* genomes is reported in Table D12 available at http://igfl.ens-lyon.fr/equipes/f.-leulier-functional-genomics-of-host-intestinal-bacteria-interactions/l-plantarum/database-for-comparative-genomics-of-54-l-plantarum-strains-7-7.

The OGs assigned to carbohydrate utilization are generally organized in “cassette”-like genomic loci, whose distributions among the strains are hypervariable and largely explain the differences found in their genome size. This finding is similar to what has been reported previously about the genomic analysis of 37 *L. paracasei* strains ^21,32^.

The initial pan-genome analysis already revealed a clustering of the 54 *L. plantarum* strains in two main groups, which appeared to include a prominent involvement of the distribution of genes belonging to sugar metabolism (Fig. 1). The hierarchical clustering of the strains based on the distribution of the OGs belonging to this category did not cluster the strains of the same source of isolation (Fig. 4). We were able to find a few weak correlations between the origin of isolation and sugar utilization genomic repertoires; 5 out of the 7 strains bearing the fewest sugar cassettes (28-31) are of human origin (NIZO1838, NIZO2256, NIZO2257, NIZO2258, NIZO3893), while 3 of the 6 strains with the most sugar cassettes (50-51) were isolated from silage (NIZO2726, NIZO2831, JDM1) (Table D1 available at http://igfl.ens-lyon.fr/equipes/f.-leulier-functional-genomics-of-host-intestinal-bacteria-interactions/l-plantarum/). Therefore, no clear correlation between sugar metabolism and niche of isolation could be found in *L. plantarum*. However, most of the sugar utilization cassettes are spread among the genomes of strains with different origins. Analogously, the allelic divergence of the genes assigned to the sugar utilization cassettes did not segragate the strains according to their origin of isolation (Supplementary Fig. 6). Overall, in contrast to the general concept proposed for several LAB species, our study based on 54 *L. plantarum* strains strongly suggest that both the presence/absence of genetic cassettes involved in sugar utilization and their allelic divergence do not reflect strain adaptation to a particular niche involving utilizing specific niche-substrates for growth. The ability of most *L. plantarum* strains to utilize many different sugars for energetic needs might explain why it is difficult to find niche determinants in this functional group.

Finally, in order to measure the degree of conservation between the phylogenetic analyses performed on these 54 *L. plantarum* genomes, we compared the phylogenies obtained for the entire core genome and for the three variomes (Fig. 5). The variome-distribution of functions assigned to the secretome and sugar metabolism functions displayed partial correlation with the core genome phylogeny, whereas the EPS/CPS-assigned variome distribution displayed very limited or no congruency with the core genome phylogeny (Fig. 5), indicating a high degree of genomic plasticity of this functional category.

**Figure 5.**
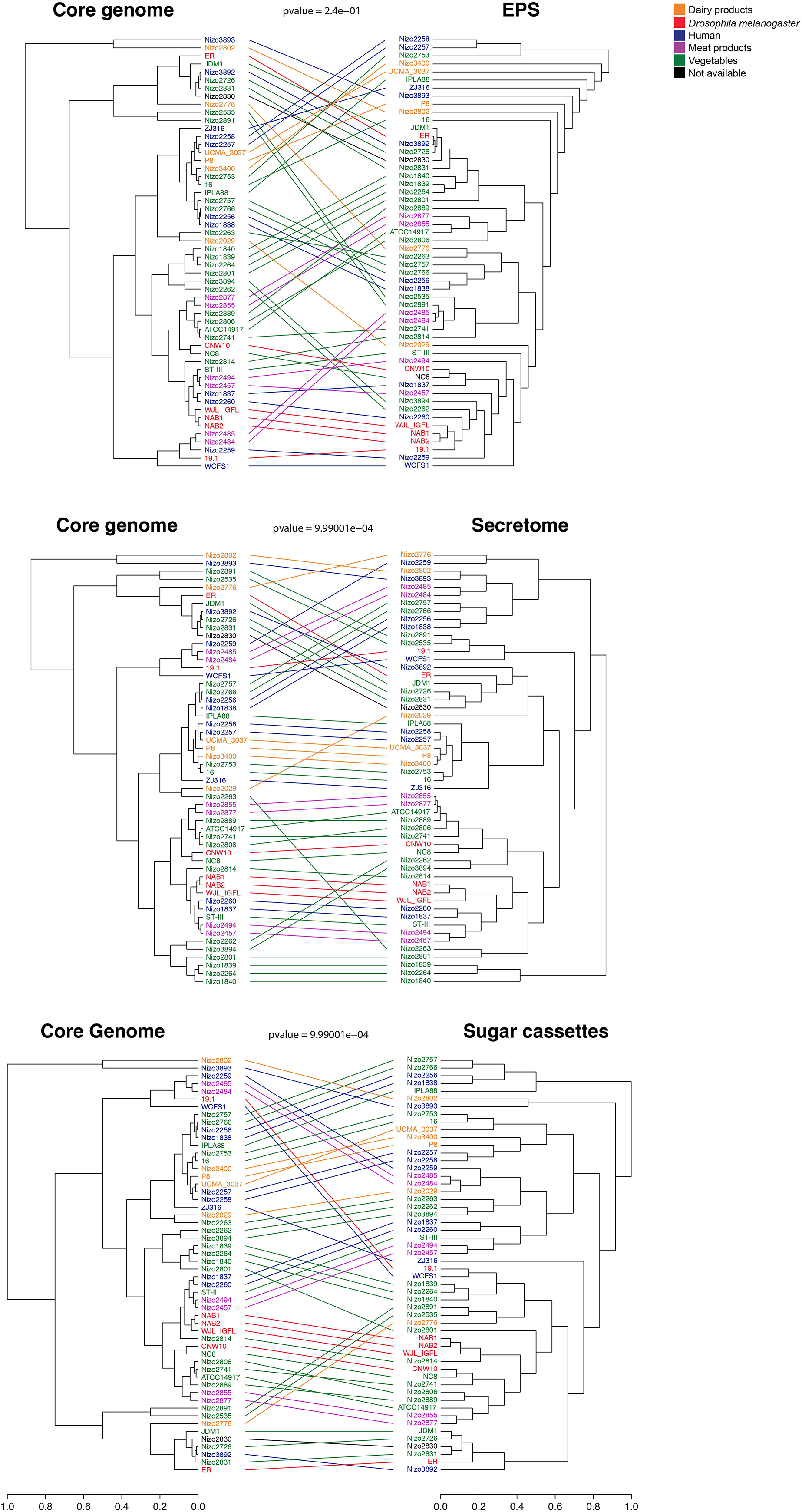
Comparison of core genome and variome phylogenies. The phylogenetic tree obtained from the analysis of the core genome has been compared to the phylogenies obtained for the 3 variome categories (EPS, Secretome, Sugar cassettes). Coloured strings connecting the same strain of both trees highlight the degree of correlation between the phylogenies. The pvalue resulted from the permutation test is displayed for each comparison.

Taken together, none of the genomic clustering approaches used in our study identified a genomic signature that reflects the origin of isolation among the of the 54 *L. plantarum* strains. The only apparent exceptions are the 7 pairs of strains that consistently appeared as close relatives in all phylogenetic analyses (NIZO2891 and NIZO2535, NIZO2484 and NIZO2484, NIZO2877 and NIZO2855, NIZO2260 and NIZO1837, NIZO2494 and NIZO2457, NIZO2831 and NIZO2726, NIZO2262 and NIZO3894) and shared their origin of isolation but geographical origins as well (Table 1). Such sharing of geographic origin prevents us from attributing a shared evolutionary history of niche adaptation to such paring.

## Discussion

We report for the first time a comprehensive sequence-based pan-genome analysis of *L. plantarum* strains isolated from different sources with the aim to further investigate their evolution and find potential genomic signatures that may reflect strain-specific niche adaptation. The processes of bacterial adaption to specific environments have been chacterized in detail^5^ ^33^. During evolution, most bacteria get rid of useless functions and enrich others that can increase fitness and survival in a particular niche ^2,15,31,32^. However, *L. plantarum* does not seem to follow this trend. Unlike most bacterial species, the evolutionary history of *L. plantarum* does not appear to be related to environmental features, such as the niche where it was isolated from. Our study suggests that *L. plantarum* is a diverse and versatile species that acquires and retains functional capacities independently of its niche, thus representing a typical example of a nomadic bacterial species. Its evolutionary history appears complex and not related to niche adaptation: its variable and flexible genetic composition helps the bacteria maintain and employ a “universal” set of genes to thrive in many different environments. Although the conventional and mainstream understanding dictates that the presence of specific gene cassettes is often encountered in bacterial adaptation to a particular niche, *L. plantarum* appears to represent an exception. We performed different analyses targeting different genomic features and parameters (GTM, allelic and functional divergence, hierarchical clustering and phylogenetic analysis, using core genome and variome separately) but failed to identify genomic signatures that reflect niche adaptation. Concerning sugar metabolism, *L. plantarum* can grow on a large variety of carbohydrates, but our results did not show an association between sugar metabolism and the strain’s origin of isolation. This result differs from what has been proposed to be a common evolutionary feature in other species of *Lactobacillus* ^2,15,31^. Strains belonging to *L. reuteri* have been shown to follow different trends of genome evolution depending on their source of isolation (i.e., rodents or humans) ^15^. Two distinct geno-phenotypes were identified in *L. rhamnosus* species by Douillard and colleagues ^31^, although a niche enrichment was confirmed only for strains isolated from the human intestinal tract by Ceapa and coworkers ^34^. Some *Lactobacillus* species isolated from the human vaginal tract, such as *L. crispatus, L. gasseri, L. jensenii* and *L. iners*, have been reported to contain smaller genomes than those of the non-vaginal species ^19^. Caffrey et al. also demonstrated fundamentally different evolutionary trends between host-associated species and their free-living relatives ^30^ One bacterial species that seems to be closer to *L. plantarum* in terms of its evolutionary relatedness and niche adaptation is *L. paracasei.* Smokvina and colleagues performed a broad comparative analysis across several *L. paracasei* genomes and also failed to identify specific evolutionary signatures related to niche adaptation, although the dairy isolates appeared to cluster together. We therefore conclude that *L. plantarum* is likely a nomadic species like *L. paracasei*, and its genomic adaptation may be driven by alternative selective pressures other than specific niche-adaptation. Nomadic lactobacilli maintain genomic flexibility that enables them to grow efficiently in a variety of environments, while the specialized lactobacilli evolved to an “evolutionary end” by specialising to a particular niche. This phenomenon might be explained by the presence of an initial common gene pool shared among the members of a bacterial species, which eventually mutates in strains that go through niche adaptation. Our findings suggest that for *L. plantarum*, the evolutionary tranjectory of the genome cannot be informatively derived from the origin of isolation, thus highlighting the potential capability of *L. plantarum* to frequently migrate across different environments. This feature probably stems from the metabolic flexibility of this bacterial species, which would buffer the selection pressure imposed by a fluctuating environment and permit *L. plantarum* to survive in variable environmental niches without accumulating massive genetic variations. At the same time, the process of niche adaptation is very complex and not strictly related to the presence of genetic markers; it might also rely on alternative regulatory mechanisms such as gene expression and protein stability.

Nevertheless, we suspect that some aspects of the approaches adopted in this study may have influenced our findings. First of all, it is always possible to encounter sampling bias in comparative genome studies. We chose to analyse *L. plantarum* strains isolated from the main environments where this species has been commonly found. However, it is important to realise that the isolation of a strain from a particular niche does not only reflect strain adaptation to that environment. Finding a strain from a given niche might be casual, and does not necessarily indicate that this niche is the environment to which the species has adapted. Also, the plant-derived isolates were not isolated from a single homogenous niche but instead from several sources which differ in terms of chemical conditions and sugar content (such as fermented fruits, different vegetables and silages). Hence, this might explain why those strains do not share a common genotype. However, we did not identify any specific signatures of niche isolation in the subgroups belonging to this category. Secondly, so far *L. plantarum* has not been demonstrated to be a host-restricted species, thus implying that the strains isolated from the animals might have another origin. In particular, we only used strains isolated from the intestinal tract for the human-derived strains. The gut is an ecosystem open to food-derived microbes. Therefore, those strains might originate from food matrices. The selection of more specific strains, such as starter cultures used in dairy and meat products and in fermented vegetables, together with strains isolated from human and animal non-gastro-instestinal sites could complement our study, and bring an added value in understanding *L. plantarum* genetics and evolution.

To conclude, this study represents a broad, extensive and in-depth comparative genome analysis of 54 *L. plantarum* strains. The analysis of the 54 *L. plantarum* genomes reveals the extreme versatility of this species. The lack of niche-specialization in these 54 strains reflects the nomadic lifestyle and discourages prolonged persistence in a specific environment, which in turn prohibits the long-term genomic adaptation and niche-specialization. Therefore, *L. plantarum* probably has acquired and retained functional capacities that enable it to effectively migrate between different niches, thus representing a prototypical example of a nomadic bacterial species that is characterized by its dynamic and flexible lifestyle.

## Methods

### *L. plantarum* isolate collection and DNA isolation

Forty-three *L. plantarum* isolates used in this study were obtained from various institutions and universities (Table 1). Eleven *L. plantarum* genomes, including *L. plantarum* reference strain WCFS1, were obtained from the National Center for Biotechnology Information (NCBI) database ^35^. All isolates were grown in standing MRS broth (Difco Laboratories, BD, US) at 37°C. Genomic DNA from each isolate was extracted using UltraClean Microbial DNA Isolation Kit (MoBio Laboratories Inc., Carlsbad, CA, USA), following manufacturer’s instructions.

### Genome sequencing and annotation

Genomes of the 38 *L. plantarum* strains obtained from the NIZO culture collection, together with the WJL_IGFL strain, were sequenced using Illumina Miseq technology, while sequencing of 4 *L. plantarum* strains (ER, NAB1, NAB2, 19.1) obtained from different institutions were accomplished using Ion Torrent PGM technology (Life Technologies). The reads of each strain were assembled into contigs using Ray ^36^ The RAST annotation server ^37^ was used to find open reading frames (ORFs) that could code for proteins and to provide an automatic annotation of the encoded functions. A RAST annotation was also done for the 5 *L. plantarum* partial genomes available in the NCBI database to allow for a straightforward comparison of annotations across all genomes. Orthologous groups (OGs) were determined using OrthoMCL ^38^. When an OG contained more than one gene per strain (i.e. paralogues), the OG was manually split into separate OGs containing only one gene per strain (except for transposase and mobile elements). The obtained contigs were aligned and ordered based on OGs using 11 published *L. plantarum* genomes (6 complete and 5 draft genomes) (Table 1) as templates. Whole genome comparisons and pseudogene analyses were completed using the Artemis Comparison Tool (ACT) ^39^ and NCBIBLAST ^40^. Metabolic pathways were analysed using KEGG pathway (http://www.genome.jp/keg/pathway.html). The CRISPRs Finder tool (http://crispr.u-psud.fr/Server/) was used to search for CRISPR direct repeats and spacers in the 54 *L. plantarum* strains.

## Plasmid prediction

Contigs that represent fragments of putative plasmids were predicted based on one or more of the following criteria: they do not map to the reference chromosomes, they encode typical plasmid functions, they map to published *L. plantarum* plasmids, they have considerably lower GC content (i.e., <40% GC) than typical *L. plantarum* chromosome (44.5%), they have at least 2x higher sequence coverage than the chromosomal contigs, they appear to be circular, they contain many mobile element proteins (transposases, recombinases, etc.).

## Gene trait matching (GTM)

In order to correlate observed phenotypes with the presence/absence of particular genes and to extend the previous analyses conducted on *L. plantarum* using CGH data ^25^, a GTM approach was performed using Phenolink, a webtool that associates bacterial phenotypes to omics data ^27^. Niche data were divided into 5 classes (vegetables, human, dairy products, meat products, not available). Phenolink performs GTM analyses for the chosen phenotype and generates a single table of OGs correlating to the classes.

## Hierarchical clustering, Phylogeny and Cluster Analysis of Functional Shifts (CAFs)

The core genome of the 54 *L. plantarum* strains consisted of OGs with gene members present across all strains was analysed using a maximum likelihood tree based on concatenated amino acids that differ between the aligned core proteins.

The hierarchical clustering was generated using Unweighted Pair Group Method with Arithmetic Mean (UPGMA) algorithm. The presence/absence of the OGs belonging to the 3 variome categories (EPS, secretome and sugar cassettes) has been used as input data. The phylogenetic trees of the 3 variome categories were generated using MEGA v5.03 ^41^. Genetic distances were computed using the Kimura two-parameter model, and the phylogenetic trees were constructed using the neighbour-joining method with translated amino-acid sequences ^42^ using the concatenated sequences of the OGs shared across the 54 *L. plantarum* strains.

Correlation tests between trees have been performed using the R package *dendextend* and its function *cor_corphenetic* that gives the cophenetic correlation coefficient for two trees. A dedicated R script has been created to perform a permutation test and obtain a pvalue (< 0.05) indicating if the given coefficient could have been obtained by chance.

The analysis of functional divergence across the 54 *L. plantarum* strains has been conducted using CAFs software ^30^ As a first step the orthologous groups were selected and a dendrogram was calculated for each gene. Consequently, functional divergence is identified as the potential departure of the derived protein function from its ancestral one as a result of amino acid changes at important functional sites ^30^. The software identifies amino acid positions within a protein, which show radical and statistically significant substitutions among clusters. This analysis was conducted on each tree by comparing the amino acid composition between two clusters to that of an outgroup using a BLOSUM62 amino acid substitution matrix ^43^ Next, the software performs enrichment tests to identify strains and categories of genes that went through significantly more (enriched) or significantly less (impoverished) functional divergence compared to a background level. The enrichment status of each category is then calculated. Finally, a heatmap is generated from the enrichment status of functional categories within strains.

## Acknowledgments

The authors are grateful to Dr Roland Siezen for his help during genomes annotations and comparative analysis, to Dr. Tom Delmont for the help in using Anvi’o software and to Dr. Dali Ma for proofreading. The work was funded by an ERC starting grant (FP7/2007-2013-N°309704). MEM was funded by the European Union’s Horizon 2020 research and innovation programme under the Marie Sklodowska-Curie grant agreement N° 659510. The lab of FL is sponsored by the EMBO YIP program, the ATIP/AVENIR program, the fondation FINOVI and the “Fondation Schlumberger pour l’Education et la Recherche”.

## Author contributions

F.L. and S.A.F.T.H conceived and directed the research. J.R.B. and M.E.M. performed the comparative genome analysis and analysed the processed data with advice from S.A.F.T.H and M.K. B.C. performed the CAFs analysis. M.W. conducted the study on the LPSM. P.J. carried out statistical analyses. S.H. and B.G. performed the Ion Torrent sequencing. M.E.M wrote the paper with inputs from all authors.

## Supplementary Information

**Supplementary information Figure 1. Pan-genome and core-genome size**. The number of pan-genome OGs (a) and core genome OGs (b) is shown as a function of genomes added to the pan-genome.

**Supplementary information Figure 2. Cluster of orthologous groups belonging to the core genome and their relative presence across the 54 *L. plantarum* strains**.

**Supplementary information Figure 3. OGs number per strain**.

**Supplementary information Figure 4. Phylogenetic tree of OGs belonging to EPS functional category**. The 54 *L. plantarum* strains were clustered based on the nucleotide sequences of the genes belonging to the EPS category. The heatmap next to the tree shows the distribution of the OGs belonging to the EPS functional category across the 54 *L. plantarum* strains. Each row corresponds to one strain and each column corresponds to one OG.

**Supplementary information Figure 5. Phylogenetic tree of OGs belonging to secretome functional category**. The 54 *L. plantarum* strains were clustered based on the nucleotide sequences of the genes belonging to the secretome category. The heatmap next to the tree shows the distribution of the OGs belonging to the secretome functional category across the 54 *L. plantarum* strains. Each row corresponds to one strain and each column corresponds to one OG.

**Supplementary information Figure 6**. Phylogenetic tree of OGs belonging to sugar metabolism functional category. The 54 *L. plantarum* strains were clustered based on the nucleotide sequences of the genes belonging to the sugar metabolism category‥ The heatmap next to the tree shows the distribution of the OGs belonging to the sugar metabolism functional category across the 54 *L. plantarum* strains. Each row corresponds to one strain and each column corresponds to one OG.

**Supplementary information Table 1. List of the 4137 novel OGs that are not present in *L. plantarum* WCFS1 reference genome**.

**Supplementary information Table 2. Gene-trait matching analysis by Random Forest classification**. Genotype-phenotype linkage analysis on origin of isolation.

**Supplementary information Table 3. List of the OGs belonging to the enriched COGs resulted from CAFs analysis**.

